# Proteomics of steady-state and culture-derived human erythropoiesis reveals the dynamics of erythroid stress

**DOI:** 10.64898/2026.10.13.758834

**Authors:** Christina Maria Perraki, Kovi Rishya Shrung, Sadaf Fazeli, Yaroslav Kaminskiy, Ann-Charlotte Björklund, Indira Barbosa, Mikaela Hillberg Widfeldt, Janne Lehtiö, Eva Hellström-Lindberg, Teresa Mortera-Blanco, Rui Mamede Branca, Pedro Luis Moura

**Author notes:** These authors have contributed equally (Christina Maria Perraki, Kovi Rishya Shrung, Sadaf Fazeli). These authors have contributed equally (Rui Mamede Branca, Pedro Luis Moura). Correspondence: Rui Mamede Branca, Department of Oncology-Pathology, Karolinska Institutet, Stockholm, Sweden. Correspondence: Pedro Luis Moura, Department of Medicine, Huddinge (MedH), Karolinska Institutet.

## Abstract

To generate a mature red blood cell, erythropoiesis requires the committed erythroid progenitor cell and later erythroblast stages to undergo a continuum of extreme morphological and functional remodeling. *In vitro* culture has been crucial to understand the extensive proteome dynamics that underlie this process. However, the millions of cells required for conventional proteomics technologies preclude the study of erythropoiesis within the steady-state human bone marrow (BM). Thus, culture-induced stress may shape our understanding of erythroid maturation. To address this knowledge gap, we developed an optimized methodology for low-cell proteomics (pauciproteomics) and pursued total proteomic analysis of paired steady-state BM and culture-derived erythroid precursor stages from 11 healthy donors. This approach defines four main protein expression patterns during BM erythroid maturation which are only partially recapitulated by cultured cells. Culture-induced stress rewires the erythroid proteome from early stages of differentiation, reshaping major transcription factor networks, kinase pathways, metabolic enzymes and erythrokine regulation. Notably, single-cell RNA sequencing analysis of steady-state healthy and myelodysplastic syndromes (MDS) BM demonstrates that the cultured cell proteome parallels gene expression dynamics within the MDS BM, modelling the regulation of *in vivo* stress erythropoiesis. Culture-specific alterations were mostly restricted to extensive carbonic anhydrase dysregulation and embryofetal globin switching, both potentially linked to altered gas exchange dynamics in culture. In conclusion, we reveal the steady-state erythroid proteome and its substantial remodeling by culture-induced stress, reframing existing knowledge on erythroid differentiation, defining new elements of stress erythropoiesis and guiding future studies in the field.

## Introduction

In steady-state conditions, the human bone marrow (BM) maintains a constant balancing act to replenish damaged and/or senescent erythrocytes, maintaining a remarkable steady-state production rate of 2 million new erythrocytes per second.^1^ This process is maintained throughout life and requires the ability to homeostatically respond to a need for increased oxygen delivery in emergency situations,^2^ e.g. due to short-term transient bleeding caused by trauma, hemolysis, dyserythropoiesis, or adaptation to environmental oxygen conditions.^3^ The group of mechanisms underlying this emergency erythroid cell production are commonly referred to as stress erythropoiesis.^4^ These are primarily regulated by the kidney-secreted endocrine hormone erythropoietin, and together with a host of downstream signaling networks, act to increase erythroid cell production and survival.^5^ On the other end of the spectrum, constant over-production of erythroid cells will eventually be damaging to the body,^6^ thus requiring fine control to avoid erythroid cell over-production after hypoxic stress resolution.

To generate a mature red blood cell, the committed erythroid progenitor cell and later erythroblast (EB) must undergo constant cell divisions accompanied by extreme phenotypical shifts, including hemoglobinization, membrane and cell surface remodeling, karyopyknosis and enucleation. Although these processes are understood to form a continuum, major studies in the field have intricately summarized erythroid differentiation as a series of discrete steps with recognizable surface immunophenotypes and morphological states.^7–10^ Because early progenitor stages are relatively infrequent in the BM as compared to the nearly-terminal orthochromatic erythroblasts, *in vitro* cell culture of hematopoietic progenitors toward the erythroid lineage has long served as a key methodology for such studies.^11^ In particular, in the modern multiomics era, proteomics of *in vitro* culture-derived erythroid precursors and terminal reticulocytes has proven a key method to understand erythroid cell differentiation.^12,13^

In spite of these advanced methodologies, little is currently known regarding the proteomic landscape of erythropoiesis within the human bone marrow (BM). The millions of cells required for conventional proteomics methods are incompatible with the analysis of defined erythroid precursor subsets, and it is presumed that their proteome is similar. Yet, the constant overabundance of erythropoietin, growth factors and nutrients within the culture setting does not recapitulate the human BM microenvironment (*e.g*. erythropoietin 100 to 1000-fold higher in culture than in blood serum, glucose 5-fold higher, *etc*.^11^). Thus, *in vitro* culture settings likely drive processes exclusive to stress erythropoiesis which we currently expect are commonplace during normal, steady-state erythroid differentiation. Further, one day of human BM erythroid cell production (∼1.7 x 10^11^ cells) easily eclipses the monthly production capacity of a well-resourced laboratory. Defining the factors that distinguish steady-state from stress erythropoiesis has implications beyond basic biology and could serve as a basis for the future improvement of erythroid-stimulating agents, as well as optimization of *ex vivo* cell production.

To address this knowledge gap, we set out to define the proteomic landscape of human steady-state and *in vitro* culture-stressed erythropoiesis. Leveraging a newly-developed and optimized method for low-cell proteomics (pauciproteomics),^14^ we investigated paired BM and culture-derived erythroid precursors from 11 healthy donors. Coupled with integrative transcriptomic analyses of stress erythropoiesis within *in vivo* myelodysplasia and *ex vivo* cell culture, we reveal the impact of culture and stress erythropoiesis on the erythroid proteome.

## Results

### Erythroid culture disconnects surface immunophenotypes from morphological states

We investigated a cohort of 11 healthy bone marrow (BM) donors with paired cryopreserved BM mononuclear cell (MNC) samples. The first sample from each pair was thawed, magnetically-enriched for CD34^+^ cells and cultured *in vitro* with serum-free, defined media for 14 days.^15,16^ At D14, the second sample was thawed, and erythroid cell subsets were purified from both samples through fluorescence-assisted cell sorting (FACS) of CD71, CD105 and Glycophorin A immunophenotypes (based on Yan *et al*.^10^). This two-week erythroid culture enabled the concurrent isolation of 8 erythroid precursor subsets, specifically Erythroid precursor [EP] stages 1, 2 and 3+4, proerythroblasts [ProE], early basophilic erythroblasts [EB], late basophilic erythroblasts [LB], polychromatic erythroblasts [Poly] and orthochromatic erythroblasts [Ortho] (**Fig. 1A**, **Data S1**). In spite of identical culture protocols and sample handling steps, 3 of 11 cultures failed to thrive and 1 of 11 could not be processed by FACS. Thus, in total, 130 samples were processed for total proteomic analysis with pauciproteomics, encompassing 11 BM donors, of which 7 had paired cultured cells (**Fig. 1B**).

**Figure 1:**
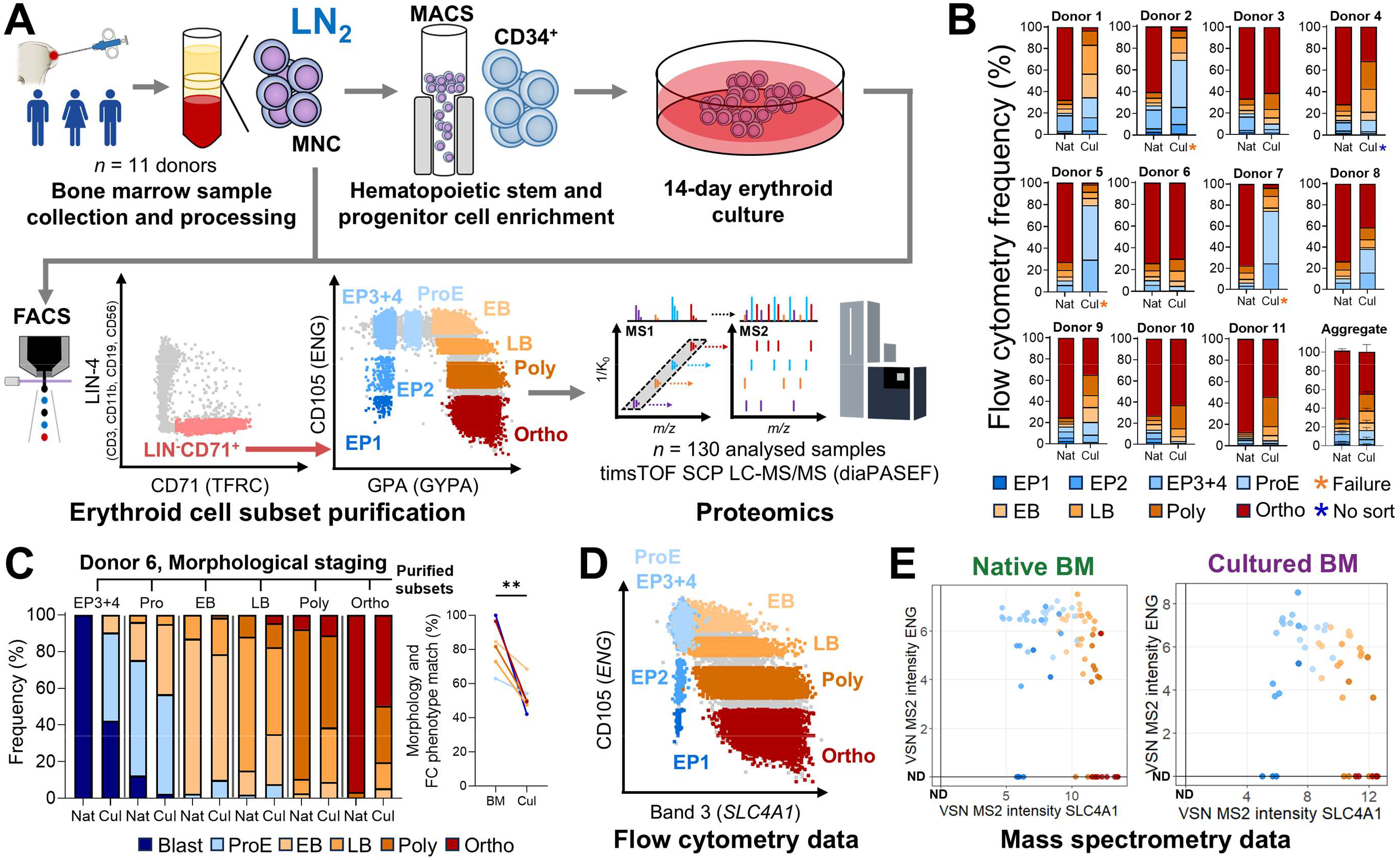
Study design and sample overview. **A)** Summary workflow for paired proteomic analysis of native bone marrow (BM)-isolated erythroid precursors (native BM) and BM cell culture-derived erythroid precursors (cultured BM). The first sample from each pair of cryopreserved (LN2) bone marrow mononuclear cells (MNC) was thawed, magnetically-enriched for CD34^+^ cells (MACS) and cultured *in vitro* with serum-free, defined media for 14 days. At D14, the second paired sample was thawed, and erythroid cell subsets were purified from both samples through fluorescence-assisted cell sorting (FACS) based on CD71/TFRC, CD105/ENG and GPA/GYPA surface expression (Yan H *et al*.^10^), totaling 130 samples across 8 erythroid cell subsets. The flow cytometry gating strategy example corresponds to native BM data from a representative donor. After isolation, samples were processed for low-cell total proteomics (pauciproteomics). **B)** Cell population frequencies quantified across donor and culture status. Orange asterisks indicate donors with culture growth failure (n = 3). **C)** Morphological staging of erythroid precursor states across FACS-purified immunophenotypes and culture status. The dot plot shows cell frequencies with successfully corresponding immunophenotype and morphology. ** = p < 0.01. **D)** Representative flow cytometry plot (native BM) displaying Band 3/SLC4A1 (X-axis) and CD105/ENG (Y-axis) expression, grouped by CD105-GPA immunophenotypes. **E)** Scatter plots of Band 3/SLC4A1 (X-axis) and CD105 (Y-axis) proteomic data (variance-stabilizing normalization [VSN]-processed MS2 intensities). Each dot represents one sample. Samples have been colored by cell population and split according to origin. As proteomic data lacks a true zero, absence of detection is indicated by ND (Not Detected). **Cell population legend: EP (1-4)**: Erythroid precursors; **ProE**: Proerythroblasts; **EB**: Early basophilic erythroblasts; **LB**: Late basophilic erythroblasts; **Poly**: Polychromatic erythroblasts; **Ortho**: Orthochromatic erythroblasts.

The FACS purification of cultured cells was based on each of the respective donor-matched native BM samples to account for biological donor variation of surface protein abundance. Morphological analysis of the entire cohort was not feasible due to cell requirements for proteomic analysis; however, Donor 6 displayed the most similar distribution between culture and BM cell states and was morphologically investigated as a representative donor. Interestingly, despite identical surface immunophenotypes, only native BM cells were representative of the intended erythroid differentiation state, whereas cultured cells displayed significant deviation toward an intermediate mix of cell compartments (**Fig. 1C**), recapitulating previously observed cell distributions from *in vitro* erythroid culture.^12^ While native BM cell subsets were generally homogeneous, culture-derived precursors also displayed significant anisocytosis with aberrant hemoglobinization patterns (**Fig. S1A/B**). Total proteomic data of surface erythroid markers were supportive of the observed immunophenotypes, confirming these features were biological rather than technical in nature (**Fig. 1D/E**). Of note, we observed that the EP1 stage was primarily comprised of blasts in native BM samples, but in culture-derived samples corresponded instead to differentiated myeloid cells (**Fig. S1A**).

### *In vivo* human erythroid differentiation is organized in functional proteomic patterns

Across all samples, we achieved a total coverage of ∼9200 quantified proteins. For detailed analysis, we first focused on defining the steady-state erythroid proteome, as this has remained a major challenge in the field. To this end, cell state transitions across each erythroid differentiation stage (i.e. pseudotime) were modelled as a time-course analysis, aiming to identify proteins which were significantly differentially expressed during erythroid differentiation or whose expression alterations were significantly modulated across time. In total, this analysis resolved four major clusters of protein expression: proteins decreasing mainly from initial differentiation stages (“rapid decrease”, *n* = 699), bimodal patterns of protein expression with peaks either in early precursors or later erythroblast stages (“early bimodal” and “late bimodal”, *n* = 836 and 412, respectively), and proteins increasing mainly with terminal erythroid differentiation (“rapid increase”, *n* = 459) (**Fig. 2A, Data S2/3**).

**Figure 2:**
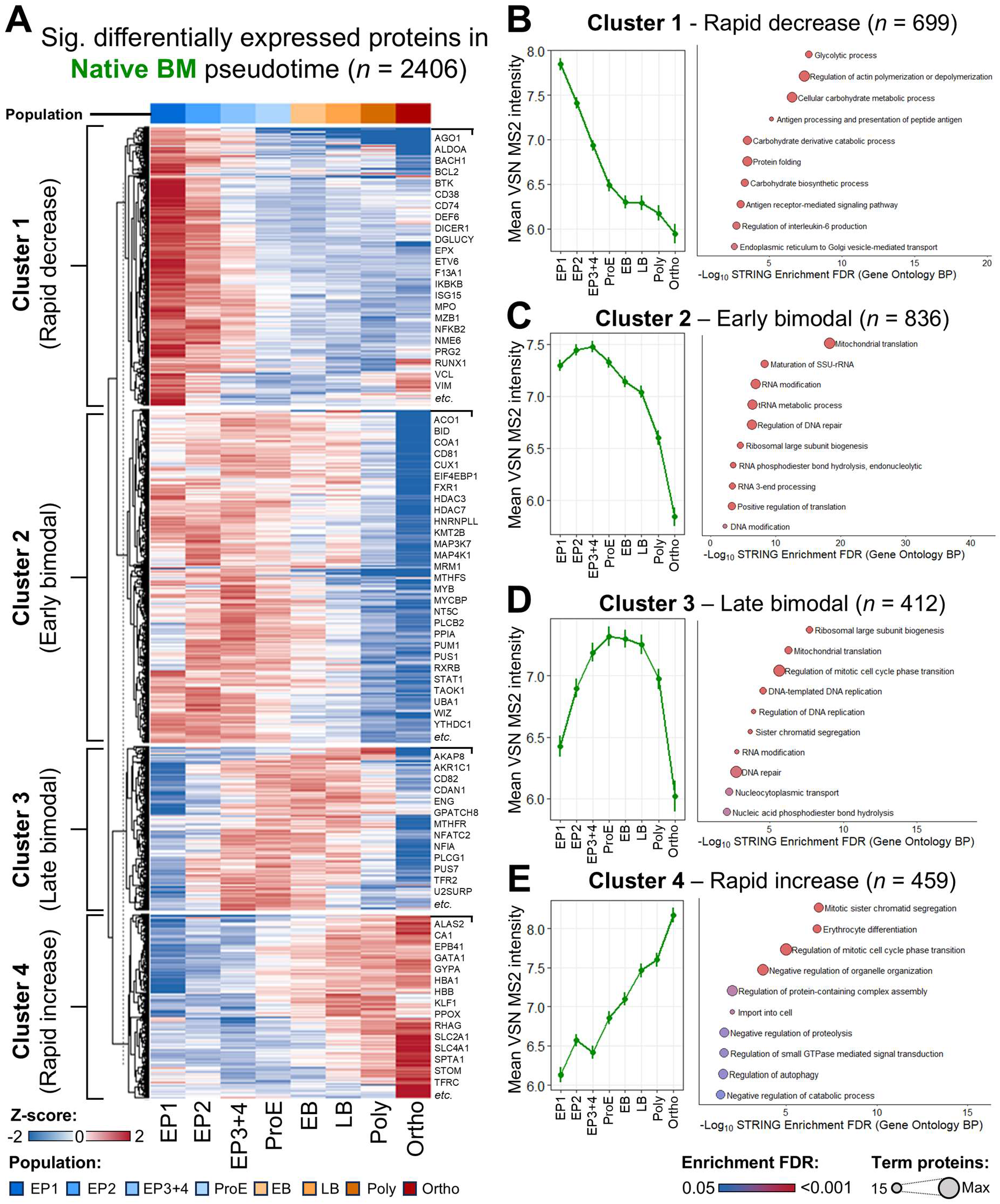
Proteome dynamics of *in vivo* human erythroid differentiation. **A)** Protein Z-score heatmap of all proteins with statistically significant differential expression (MaSigPro α = 0.10) across the pseudotime continuum of bone marrow-purified erythroid precursors. Clusters were obtained by unsupervised K-means clustering (K = 4) and labelled according to the predominant pseudotime dynamics of protein expression within the cluster. Selected example proteins for each cluster are labelled to the right. **B-E)** Mean (± standard error of the mean) expression of all proteins within a cluster (left) and STRING enrichment analysis results (right, Gene Ontology Biological Processes module) for all proteins included in **B)** Cluster 1 (Rapid decrease, n = 699), **C)** Cluster 2 (Early bimodal, n = 836), **D)** Cluster 3 (Late bimodal, n = 412) and **E)** Cluster 4 (Rapid increase, n = 459). **FDR**: False Discovery Rate.

These four proteomic patterns were associated with key functional pathways, as determined by enrichment analysis (**Data S4**). Specifically, rapidly decreasing proteins were primarily associated with non-erythroid cell identity and substantially enriched in immune proteins as well as glycan metabolism (**Fig. 2B**). Early and late bimodal proteins were associated with substantial upregulation of ribosomal biogenesis, RNA modifier enzymes and cell cycle mediators, likely linked to a need for fine translational control during the erythroid differentiation program (**Fig. 2C/D**). Finally, rapidly increasing proteins largely encompassed well-known elements for erythroid function and differentiation (**Fig. 2E**). These included transcription factors (KLF1, GATA1), membrane complexes (Band 3, Band 4.1, Glycophorins, RHAG, GLUT1/SLC2A1, Stomatin, *etc*.), iron metabolism mediators (CD71/TFRC, ALAS2, ALAD, PPOX, UROD), gas exchangers (globins, carbonic anhydrases), cytoskeletal components (ankyrin, spectrins, adducins, talin, tropomodulin, *etc*.) and additional significant enrichment of proteins involved in chromatin regulation and autophagy. These latter proteins are likely involved in karyopyknosis and enucleation (including aurora kinases AURKA/AURKB, kinesins, septins and condensins, among others), as well as later reticulocyte maturation.

### Culture-induced stress significantly remodels the erythroid proteome

Mapping the native erythroid continuum into a pseudotime landscape allows for culture-induced erythropoiesis to be directly compared. Remarkably, dimensional reduction and unsupervised clustering analysis separated the two sample groups by source upon terminal erythroid differentiation (**Fig. 3A**), despite near-identical expression patterns of key erythroid staging proteins across erythroid differentiation in both native BM and culture (**Fig. 3B**).

**Figure 3:**
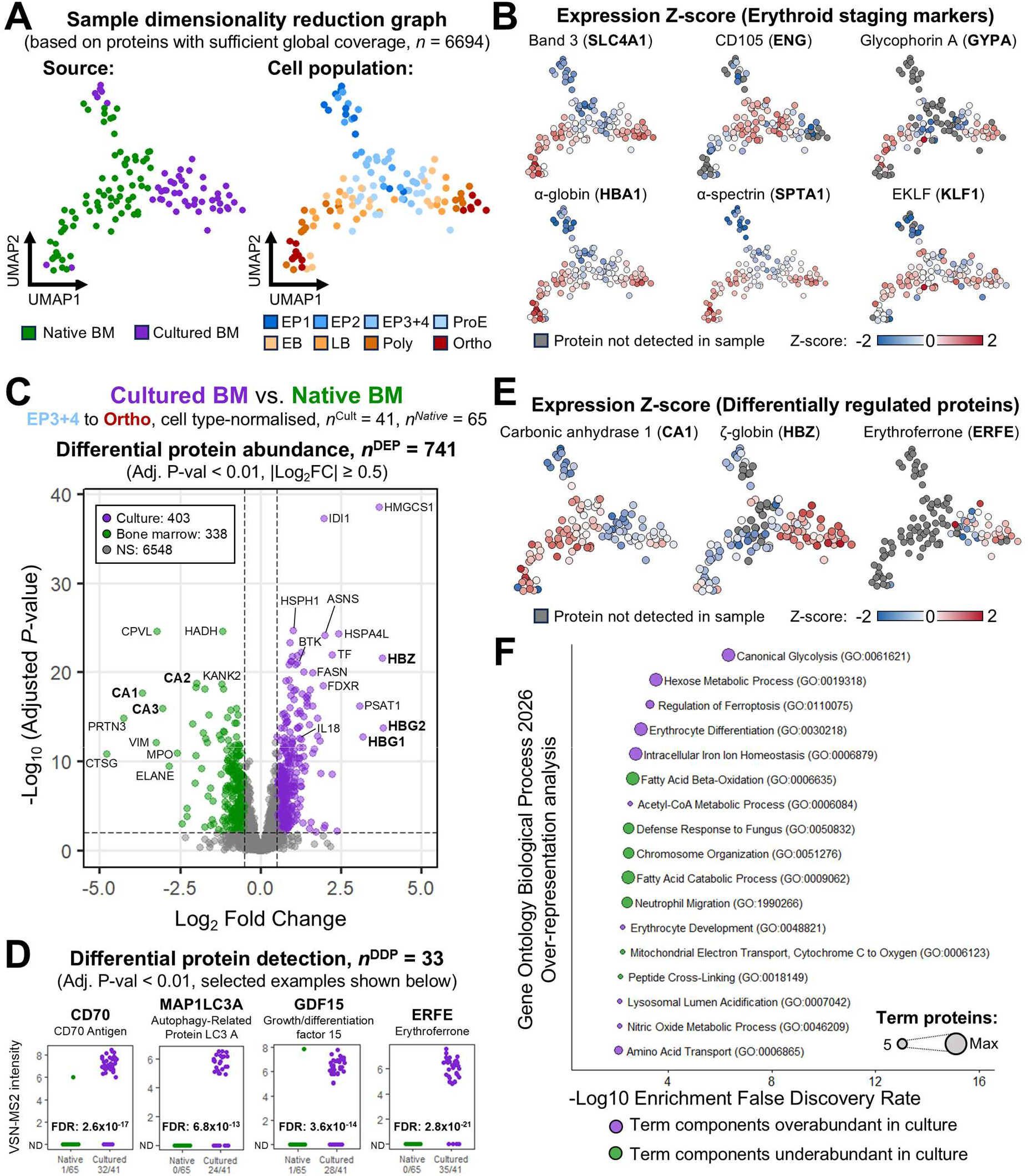
Culture-induced stress erythropoiesis disturbs key proteins for erythroid cell function. **A)** Uniform Manifold Approximation And Projection (UMAP) dimensionality reduction graph of total proteome data. Each dot represents one sample. Samples have been colored by source (left, Bone marrow-purified cells: green; Culture-derived cells: purple) and cell population (right). **B)** UMAP overlay of protein expression Z-scores for key erythroid staging marker proteins. Samples are colored by protein Z-score (from blue, to white, to red), or left grey if the protein could not be quantified in the sample. **C)** Volcano plot of differentially expressed proteins (Unpaired *t*-test Adj. P-value < 0.01, |Log2 Fold Change| ≥ 0.5, n^DEP^ = 741) between culture and bone marrow samples, encompassing all stages from EP3+4 to Ortho and normalized across cell types. Adj. P-val and Log2FC cut-offs are indicated as dashed lines. Purple and green-labelled proteins are respectively over- and underabundant in the culture setting. **D)** Dot plots of VSN-MS2 intensity for representative differentially detected proteins (Fisher’s Exact Test Adj. P-value < 0.01, n^DDP^ = 33) between culture and bone marrow samples. Each dot represents one sample, all samples have been colored by source. **E)** UMAP overlay of protein expression Z-scores for example differentially regulated proteins between culture and bone marrow. **F)** Over-representation analysis results (Gene Ontology Biological Processes 2026 module) for all differentially expressed/detected proteins between culture and bone marrow cells. For directionality purposes, terms are only included if all proteins in the term are over- (purple) or underabundant (green) in the culture setting.

Thus, we focused on comparative analysis of culture-derived cells against steady-state BM. Since EP1+2 were morphologically distinct cells between culture and native BM (**Fig. S1A**), we focused only on EP3+4 to Ortho stages for this comparison. We hypothesized that protein expression could be restricted to either native or cultured BM groups; therefore, we pursued an integrated statistical analysis to identify differentially expressed proteins (DEP) as well as differentially detected proteins (DDP) (**Data S5**). Indeed, from EP3+4 stages onward, culture-induced stress erythropoiesis was associated with widespread proteomic dysregulation. This was the case both for quantifiably differentially abundant proteins (**Fig. 3C**, n^DEP^ = 741) as well as for sample group-specific protein expression (**Fig. 3D, Fig. S2**, n^DDP^ = 33).

Notably, the culture setting was associated with highly significant overexpression of embryoid/fetal globin chains in an apparent culture-induced globin switch, despite no fetal bovine serum or similar products being included in erythroid culture media (Zeta and Gamma 1+2, Mean Log_2_ Fold Change [mLog_2_FC] = 3.58, **Fig. S3**). Furthermore, significant deficiency of all carbonic anhydrases was observed (CA1, CA2 and CA3, mLog_2_FC = -2.91) (**Fig. 3C/E**). This was accompanied by the culture-exclusive production of stress erythrokines Growth Differentiation Factor 15 (GDF15) and erythroferrone (ERFE); an extreme metabolic restructuring, in particular of cholesterol, fatty acid beta-oxidation and iron metabolism genes; and the remodeling of kinase-driven signaling pathways, e.g. STAT3 and BTK (**Fig. 3E/F**).

### Cultured erythroid cell proteomes recapitulate *in vivo* stress erythropoiesis

Whilst culture-induced dysregulation of the erythroid proteome is evident, it is unclear from these data alone whether the identified mechanisms arise from *bonafide* stress erythropoiesis or are instead an artifact of cell culture (or both). Resolving the cause of these alterations is therefore crucial to define how erythroid stress influences steady state-erythropoiesis.

To this end, we pursued an orthogonal analysis of external single-cell RNA sequencing (scRNAseq) data, integrating our previously acquired data on healthy and myelodysplastic syndrome (MDS) erythroid differentiation^15^ together with data from Berastegui *et al*.^17^ and from Triana *et al*.^18^ (strategy summarized in **Fig. 4A**). Our previous data consists of full transcriptome coverage of healthy stem and progenitor cells, as well as GPA^+^-defined erythroid precursors. The Berastegui *et al*. dataset consists of full transcriptome coverage of healthy BM CD34^+^-derived erythroid cells, in a similar setting to the cultures pursued in this study. Finally, Triana *et al*.^18^ report a relatively limited transcriptome coverage of healthy BM cells with highly detailed immunophenotypic data from antibody-derived tag analysis. We thus constructed a phenotype-informed transcriptomic atlas of healthy, MDS-stressed and culture-stressed erythroid differentiation (**Fig. 4B/C**). Importantly, the marker genes of predicted scRNAseq phenotypes matched our proteomic dataset, confirming the successful integration of these different data modalities (**Fig. 4D, Data S6**). After this step, the data from Triana *et al*. were removed from the dataset, to preserve only Chromium data with full transcriptome coverage.

**Figure 4:**
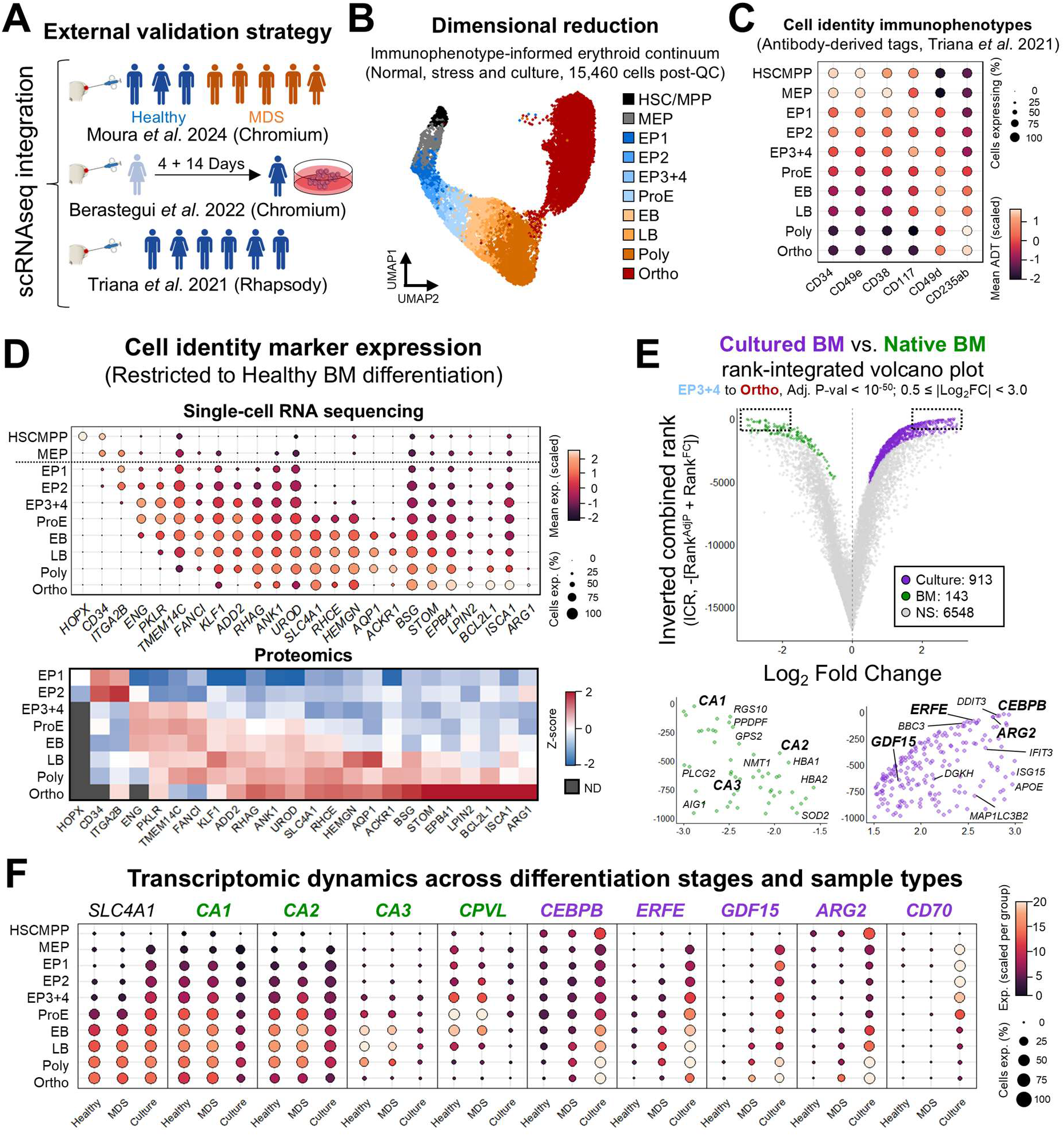
Stress erythropoiesis is similarly regulated in healthy erythroid cultures and *in vivo* myelodysplastic erythropoiesis. **A)** Strategy for data integration of single-cell RNA sequencing (scRNAseq) datasets encompassing healthy and myelodysplastic (MDS) erythropoiesis (Moura et al. 2024, bone marrow samples from 3 healthy individuals and 5 MDS patients, 10X Chromium), culture-induced stress erythropoiesis (Berastegui et al. 2022, bone marrow-isolated CD34^+^ cells were transduced with a control shRNA virus, expanded for 4 days and subjected to erythroid culture conditions for 14 days), and phenotype-informed healthy erythropoiesis (Triana et al. 2021, 6 healthy individuals, BD Rhapsody). **B)** UMAP bidimensional projection of cells integrated from the three experiments, colored according to mapped immunophenotypes. Each dot corresponds to one cell. **C)** Dotplot of antibody-derived tag marker data relevant to erythroid differentiation (from Triana *et al*.), separated by this study’s immunophenotype labelling. **D)** Cell population marker gene expression during healthy erythropoiesis across integrated scRNAseq data (top, dotplot of expression level and frequency of expressing cells) and proteomics (bottom, Z-score heatmap). **E)** Rank-integrated volcano plot of differentially expressed genes between culture-derived and bone marrow cells (Adj. P-value < 10^-50^, 0.5 ≤ |Log2 Fold Change| < 3.0, n^DEP^ = 1056). Top over- and under-expressed genes are shown below (zoom to inverted combined rank > -1000). **F)** Dotplot of comparative transcriptomic dynamics for example differentially expressed genes across healthy bone marrow erythropoiesis (Healthy), MDS bone marrow erythropoiesis (MDS) and healthy culture-derived erythropoiesis (Culture). Expression frequency for each population/group is displayed as dot size. Band 3 (SLC4A1) is shown as a control gene. Genes were labelled according to differential protein expression data shown in Figure 3 (CA1, CA2, CA3, CEBPB, CPVL = differentially expressed proteins; ERFE, GDF15, ARG2, CD70 = differentially detected proteins). **Cell population legend: HSCMPP**: Hematopoietic stem cells and multipotent progenitors. **MEP**: Megakaryocyte-erythroid progenitors. **EP (1-4)**: Erythroid precursors; **ProE**: Proerythroblasts; **EB**: Early basophilic erythroblasts; **LB**: Late basophilic erythroblasts; **Poly**: Polychromatic erythroblasts; **Ortho**: Orthochromatic erythroblasts.

Although state-of-the-art methods for scRNAseq data analysis are based on biological replicate pseudobulking, this approach was not possible, since only one culture-derived sample was included. Therefore, we pursued a heavily constrained statistical analysis of all culture-derived cells against healthy and MDS BM cells (Adj. P-value < 10^-50^; 0.5 ≤ Log_2_ Fold Change < 3.0; as before, limited from EP3+4 onward, **Data S7**). This displayed similar results to our proteomic analysis, as expected from the culture setting (**Fig. 4E**). However, we could verify that MDS samples displayed an intermediate phenotype with clear representation of the same culture-induced genes (**Fig. 4F**). Most culture effects resulted from *bonafide* erythroid stress rather than culture-specific artifacts alone, including even metabolic restructuring, albeit with less extreme effect as compared to protein data. Key exceptions confirmed as culture-specific events were deficiencies in carbonic anhydrase and *CPVL* expression, as well as the extreme upregulation of *CD70* in erythroid precursors (differentially detected culture-specific protein). Embryofetal globins displayed strange expression patterns overall, perhaps due to incorrect alignment of Chromium short reads against homologous gene regions, and their differential expression was therefore not confirmed in scRNAseq data.

### Integrated proteotranscriptomics identifies key signaling pathways for erythroid stress

The stress-related transcription factors *ATF4*, *DDIT3*, *CEBPB* and *XBP1* were across the most highly upregulated genes induced by culture. Importantly, all four transcription factors were also significantly upregulated within the MDS setting, highlighting these as potential regulators of stress erythropoiesis (**Fig. 5A**). Whilst only CEBPB could be quantified through proteomics, its level was significantly increased in culture. Specifically, CEBPB displayed increased expression in early precursors and maintained expression in late erythroblast stages, in contrast to its absence in native BM cells (**Fig. 5B**).

**Figure 5:**
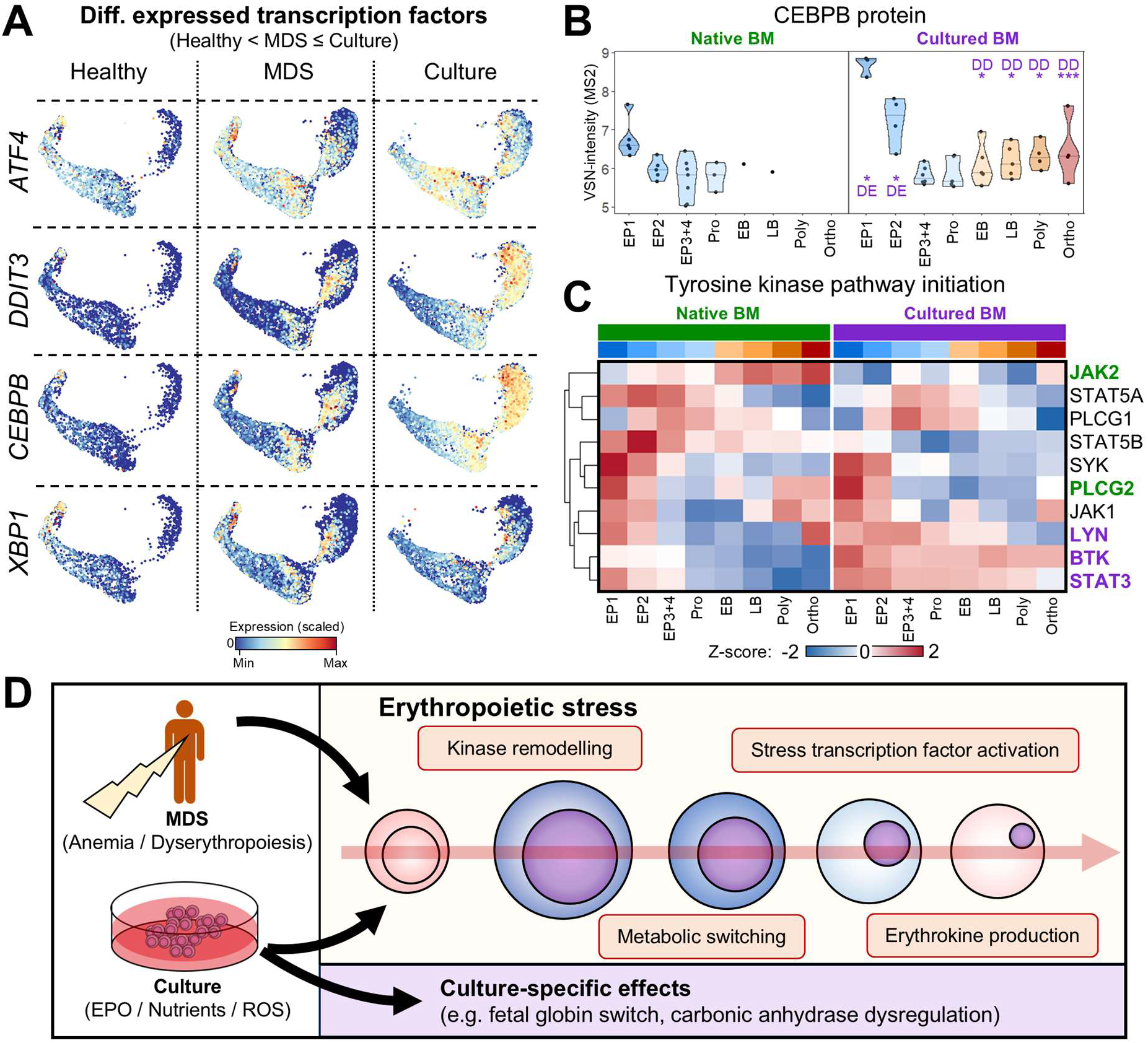
Stress erythropoiesis remodels the signaling landscape of developing erythroid cells. **A)** UMAP overlay of scaled RNA expression for differentially expressed transcription factors with gradual change across healthy, MDS and culture settings. **B)** Violin plot of VSN-MS2 protein intensity for the CEBPB transcription factor across bone marrow (left) and culture-derived (right) erythropoiesis. Each dot represents one sample, all samples have been colored by source. Differential expression and detection information are labelled per cell population. * = Padj < 0.10, ** = Padj < 0.01, *** = Padj < 0.001. **C)** Protein Z-score heatmap of signaling proteins involved in tyrosine kinase pathway initiation. Protein names have been colored according to differential expression information (black = no significance, purple = significantly overabundant in culture, green = significantly underabundant in culture). **D)** Summary diagram. MDS and culture stress induce substantial remodeling of molecular and cell biology within the developing erythroid precursor up until the end of erythropoiesis, with key features distinguishable from healthy steady-state erythropoiesis, such as kinase remodeling, metabolic switching, stress-induced transcription factor activation and erythrokine production. In addition, culture conditions act as an additional modulator of erythroid biology with particular impact on gas exchange-related proteins, exemplified by fetal globin switching and widespread dysregulation of carbonic anhydrase levels.

Since these transcription factors are regulated by upstream kinases,^19^ we evaluated overall kinase levels. We searched for regulatory mechanisms that could justify these transcription factor dynamics by focusing on kinase-mediated signaling. Culture induced a significant reduction of JAK2 concomitantly with increased levels of STAT3, LYN and BTK, showing that stress erythropoiesis completely remodels the abundance of effector kinases (**Fig. 5C**).

In summary, this paired proteomic analysis of *in vivo* and culture-derived stress erythropoiesis bypasses individual donor variation to define the impact of culture stress on the erythroid proteome. By coupling integrative transcriptomic analyses of *in vivo* MDS-associated and *ex vivo* culture-induced stress erythropoiesis, we demonstrate that most culture-induced effects are indeed key elements of human *in vivo* stress erythropoiesis. These elements include metabolic switching, kinase remodeling, activation of transcription factors not conventionally associated with erythroid differentiation, and stress erythrokine production. We also identify culture-specific effects, primarily the disruption of gas exchange proteins with carbonic anhydrase dysregulation and embryofetal globin switching (**Fig. 5D**).

## Discussion

Over decades, functional studies of human erythropoiesis have been largely informed and enabled by the ability of *in vitro* culture systems to recapitulate the full continuum of terminal erythroid differentiation from undifferentiated progenitors to terminal anucleate reticulocytes.^11^ However, across an extensive repertoire of culture conditions and cell models,^20–22^ no practical model for steady-state human differentiation exists, leaving this process as an open question. In this study, utilizing low-cell proteomics and integrative single-cell transcriptomic analyses, we pursued the first comparative proteotranscriptomic analysis of human steady-state BM erythropoiesis and BM-derived culture stress erythropoiesis. These data reveal across all erythroid differentiation states that stress erythropoiesis dynamically rewires the global transcriptome and proteome, informing basic erythroid biology and providing context for the future improvement of culture conditions as well as erythroid-stimulating therapies.^23^

Focusing on the key point of similarity, it is evident that cultured erythroid cells do generally model the proteomic shifts of steady-state erythroid differentiation. In particular, the gradual erythropoietic remodeling of cytoskeletal/membrane proteins appears to be particularly well-maintained in both total proteomics and surface immunophenotypes, suggesting that this is a carefully controlled process and likely independent from stress erythropoiesis. However, we identify striking differences between steady-state and culture-induced stress erythropoiesis already in early precursor stages which only become more pronounced during terminal erythroid differentiation. We expected this to be the case, given the significantly different energy and growth factor status within erythroid culture; however, we were surprised to find that most of these pathways were required by *in vivo* stress erythropoiesis in MDS.

These proteomic data shed new light on many existing studies. For example, stress-specific elements included ARG2 (Mitochondrial Arginase 2), recently described as a key factor in culture and Diamond-Blackfan Anemia^24^ yet here absent in the steady-state BM. Along the same lines, the extreme erythrokine production of GDF15 and ERFE, described in culture conditions^12^ and MDS^15,25^, was again absent in steady-state. Furthermore, we describe an extensive remodeling of stress transcription factor and kinase signaling, identifying the overactivation of CEBPB as a yet undescribed signaling element in late stress erythropoiesis.

It is attractive to speculate that these and other identified processes are central to all settings of stress erythropoiesis; however, further studies of the human BM focusing on different stressors will inform how erythroid production may be further modulated, including *e*.*g*. ribosomopathies and in particular Diamond-Blackfan Anemia^24^ (lack of erythroid production), aplastic anemia^26^ (BM failure), hemolytic anemia^27^ (erythroid destruction), enzymopathies and hemoglobinopathies^28^ (erythroid dysfunction). The major culture-specific finding which was not recapitulated by *in vivo* stress erythropoiesis was the extensive dysregulation of carbonic anhydrases and accompanying embryofetal globin switch. Since both protein groups are key for gas exchange,^29,30^ we hypothesize that hyperoxia and constant carbon dioxide levels may drive these alterations, which may influence the function of *ex vivo*-produced cells.

In summary, we reveal how stress erythropoiesis and *ex vivo* culture affect the developing erythroid cell. These data provide an important benchmark for future studies, defining which elements of erythroid differentiation are crucial in any context versus those that are enacted during erythroid stress, with translational relevance to studies of erythroid pathogenesis.

## Methods

### Healthy donor sample processing and ethical approval

Healthy donor samples were biobanked according to standard operating protocols at the Karolinska Institutet Biobank, Stockholm, Sweden (*n* = 11 healthy donors). In short, bone marrow (BM) samples were separated by Lymphoprep™ (STEMCELL Technologies) density gradient centrifugation at 400g for 30 min, room temperature (RT), to derive a mononuclear cell layer (MNC). MNC were cryopreserved in 50% RPMI 1640 Glutamax (ThermoFisher), 40% inactivated fetal bovine serum (FBS, ThermoFisher) and 10% Dimethyl Sulfoxide (DMSO) (Sigma-Aldrich). Cryopreserved MNC were thawed in RPMI 1640 Glutamax + 20% FBS + 100 U/mL DNase I (Sigma-Aldrich). All source material was provided with written informed consent for research use, given in accordance with the Declaration of Helsinki, and the study was approved by the Ethics Research Committee at Karolinska Institutet (2017/1090-31/4, 2022-03406-02, 2024-03119-02).

### Erythroid cell culture

BM MNC were thawed and processed for magnetic-activated cell sorting (MACS) using CD34 MicroBeads (Miltenyi Biotec) and following the manufacturer’s protocol. After separation, cells were seeded and cultured in a defined medium of IMDM (Gibco) supplemented with 15% BIT9500 (StemCell Technologies), 1% Penicillin-Streptomycin (Invitrogen), 50 ng/mL human stem cell factor (Invitrogen), 20 ng/mL interleukin-3 (Invitrogen) and 20 ng/mL interleukin-6 (Invitrogen). Media changes were performed on days 2, 4, 7, 9 and 11. On days 7, 9 and 11, culture media were additionally supplemented with 2 IU/mL recombinant human erythropoietin (PBL Assay Science) and 800 ng/mL human holo-transferrin (Sigma-Aldrich). Cells were kept in standard human cell culture conditions of 5% CO_2_ and 37°C throughout the 14-day period.

### Morphological evaluation and microscopy analysis

Cells were cytospun, fixed in methanol for 15 min at RT and air-dried. Slides were stained with May-Grünwald Giemsa using standard pathology procedures. Micrographs were acquired using a Pannoramic MIDI II slide scanner (3D Histech) at 40x with a Hitachi HV-F22 3CCD SXGA camera (Hitachi Kokusai Electric).

### Sample processing for low-cell proteomics (Pauciproteomics)

We have recently reported the development of pauciproteomics,^14^ here further optimized. Briefly, the first sample of each pair was thawed and cultured as detailed above. On day 14, the second sample was thawed, and both samples were processed for fluorescence-activated cell sorting (FACS) using an FACS ARIA II Fusion (Becton Dickinson) or a FACSDiscover™ S8 (Becton Dickinson) at the MedH FACS facility of Karolinska Institutet. All steps were performed in FACS buffer (PBS + 2% FBS + 1 mM EDTA) kept at 4°C. All experiments included fluorescent-minus-one (FMO) and single-stained controls. Antibodies used are listed in **Table S1**. Cells were FACS-purified onto the center of dry Protein LoBind tubes (Eppendorf) (sorting strategy in **Fig. 1A**, based on Yan *et al*.^10^). After sorting of up to a maximum of 1000 cells, samples were snap frozen in dry ice and later prepared for mass spectrometry analysis using a procedure of digestion on Evotip based on and modified from Ye *et al*.^31^. Finally, samples were analyzed for LC-MS/MS using a Evosep One (Evosep) coupled to a timsTOF SCP mass spectrometer (Bruker) in data independent acquisition diaPASEF (parallel accumulation-serial fragmentation) mode. Extended details are provided in **Sup. Methods**.

### Data analysis and statistical methods

Proteomics: Raw mass spectrometry data files were analyzed using Spectronaut v.19.4 (Biognosys) with the DirectDIA workflow for label-free quantification. Resulting MS2 protein intensity matrices were analyzed with a customized pipeline for combined statistical analysis of differential expression (FDR-adjusted unpaired t-test), differential detection (FDR-adjusted Fisher’s Exact Test) and time-course analysis (two-regression step approach from MaSigPro^32^).

Single-cell RNA sequencing: Data matrices were obtained from Moura *et al*.^15^ (10X Chromium, RNA data from BM samples of 3 healthy individuals and 5 myelodysplastic syndromes [MDS] patients), Berastegui *et al*.^17^ (10X Chromium, RNA data from BM-derived control transduced CD34^+^ cells after 4 days of expansion and 14 days of erythroid culture conditions) and Triana *et al*.^18^ (BD Rhapsody, RNA and antibody-derived tag [ADT] data from BM samples of 6 healthy individuals). Data were processed, integrated and analyzed using Seurat v.5.4.0.^33^ Extended details for high-throughput data analysis are provided in **Sup. Methods**.

Gene set enrichment and over-representation analyses were performed via the STRING database (v. 12.0).^34^ All statistical analyses were performed with RStudio v.2026.07.1.147, R v.4.5.3 and GraphPad Prism v.11.0.0. Flow cytometry data were analyzed using FlowJo v. 10.10.0 (Becton Dickinson). Image analysis and cell area quantification was performed through manual annotation and automated area calculation using QuPath v. 0.7.0.^35^

## Acknowledgments

The authors would like to thank the healthy donors for their willingness to participate in this research, as well as the crucial biobanking activities and sample access provided by the Karolinska Institutet Biobank (Stockholm, Sweden). Mass spectrometry analyses were performed using instrumentation located at the Clinical Proteomics Mass Spectrometry facility, Karolinska Institutet / Karolinska University Hospital/ Science for Life Laboratory. The authors also thank Dr. Timothy Satchwell (University of the West of England, Bristol, UK) for productive scientific discussions.

CMP was supported by a Karolinska Institutet Doctoral funding grant (Dnr 2025-01553). KRS, YK, JL and RMB were supported by Vetenskapsrådet (grant number 2021-00754), Erling Persson Stiftelse (grant number 22/9-2020), Cancerfonden (grant numbers 20 1269 Pj F, 23 2819 Pj 01 H), VINNOVA (2024-01137), the Cancer Research Foundations of Radiumhemmet (grant number 241272) and the Stockholm County Council (SLL) Region Stockholm (grant number FoUI-1000396). EHL was supported by Cancerfonden (grant number 19 0200), Vetenskapsrådet (grant number 211133), Knut and Alice Wallenberg Foundation (grant number 2017.0359). PLM was supported by the European Hematology Association (Advanced Research Grant Topics-in-Focus Precision Hematology), the Dr. Åke Olsson foundation (Dnr 2024-00303), the Alex and Eva Wallström foundation (Dnr 2024-00311), the Jeanssons Stiftelser (J2025-0123), the Felix Mindus contribution to Leukemia Research (Dnr 2025-02863) and a KI Research Foundation grant (Dnr 2024-02330).

## Authorship contributions

CMP and PLM conceived and designed the study. CMP, KRS, SF, JL, EHL, TM-B, RMB and PLM contributed to experiment design and result interpretation. CMP, SF, TM-B and PLM conducted cell culture and FACS experiments. KRS, YK and RMB conducted mass spectrometry experiments. A-CB, IB, MHW and EHL conducted or administered clinical biobanking activities. CMP, SF and PLM conducted computational analyses. CMP, KRS, SF, RMB and PLM created figures and wrote the manuscript. All authors read, edited, and approved the manuscript.

## Data availability

The mass spectrometry proteomics data (including raw files, search files, search parameters and fasta file) have been deposited to the ProteomeXchange Consortium via the PRIDE partner repository with the dataset identifier **PXD083653** (*Low-cell Proteomics of bone marrow-collected and culture-derived human erythroid cells*). Other raw data are available from the corresponding author upon request.

## MAIN SUPPLEMENTAL FILE

### Supplemental data descriptions

**Data S1**: Map of low-cell samples collected for erythroid cell pauciproteomics.

**Data S2**: MaSigPro analysis of differentially regulated proteins across steady-state erythroid differentiation.

**Data S3**: List of proteins within differentially expressed protein clusters of progression through steady-state erythroid differentiation.

**Data S4**: STRING enrichment analysis of functional proteomic enrichments in steady-state erythroid differentiation clusters.

**Data S5**: Differential protein expression between culture-derived and steady-state erythropoiesis.

**Data S6**: Cell type-specific RNA expression profiles in integrated data of steady-state, *in vivo* stress (MDS) and *ex vivo* stress (culture) erythropoiesis.

**Data S7**: Differential RNA expression between culture-derived and bone marrow erythropoiesis (combined healthy and MDS).

## Supplemental figures

**Figure S1:**
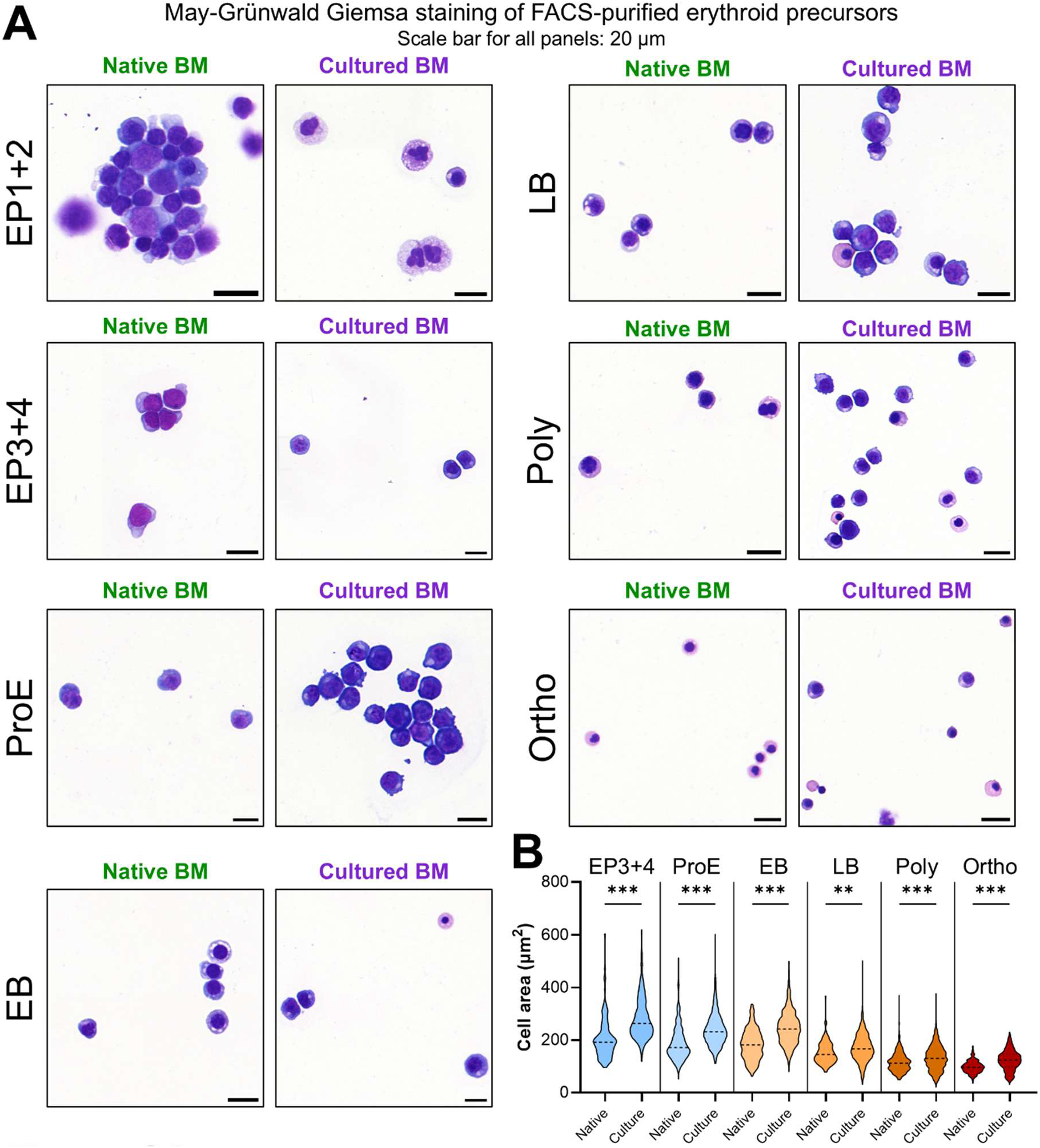
Morphological analysis of native and culture-derived bone marrow erythropoiesis. **A)** Micrographs of FACS-purified erythroid cell populations (gating strategy in Figure 1). Erythroid precursors 1+2 were purified together in these data. After FACS-purification, cells were cytospun, methanol-fixed and stained with May-Grünwald Giemsa. Scale bars = 20 μm. **B)** Cell area of erythroid cell populations separated by sample source. Adjusted P-values correspond to Holm-Sidak correction of unpaired T-tests between each source, within each population. **Cell population legend: EP (1-4)**: Erythroid precursors; **ProE**: Proerythroblasts; **EB**: Early basophilic erythroblasts; **LB**: Late basophilic erythroblasts; **Poly**: Polychromatic erythroblasts; **Ortho**: Orthochromatic erythroblasts.

**Figure S2:**
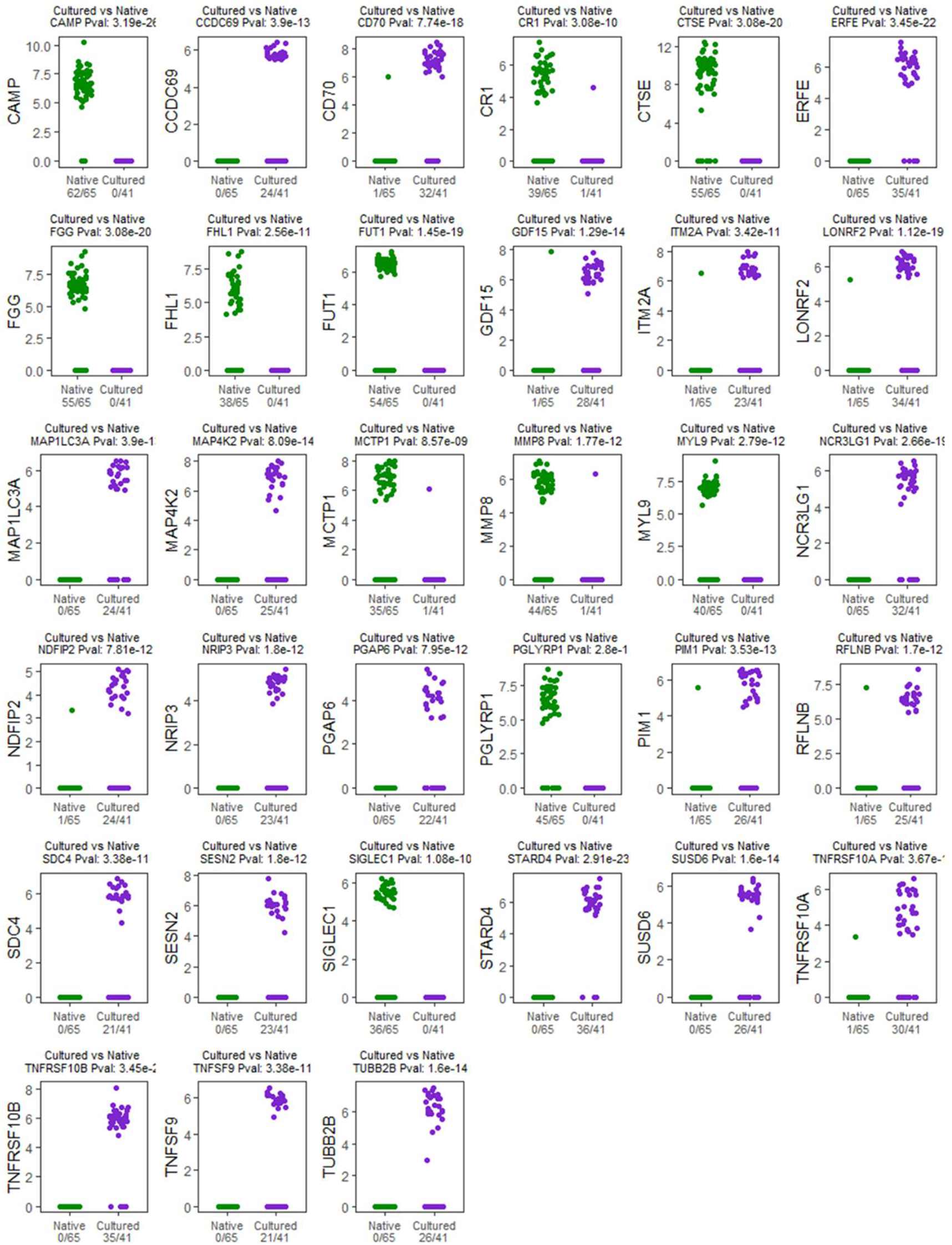
Differentially detected proteins between culture-derived and *in vivo* BM erythropoiesis. Dot plots of VSN-MS2 intensity for differentially detected proteins (Fisher’s Exact Test Adj. P-value < 0.01, n^DDP^ = 33) between culture and native BM samples. Each dot represents one sample, all samples have been colored by source.

**Figure S3:**
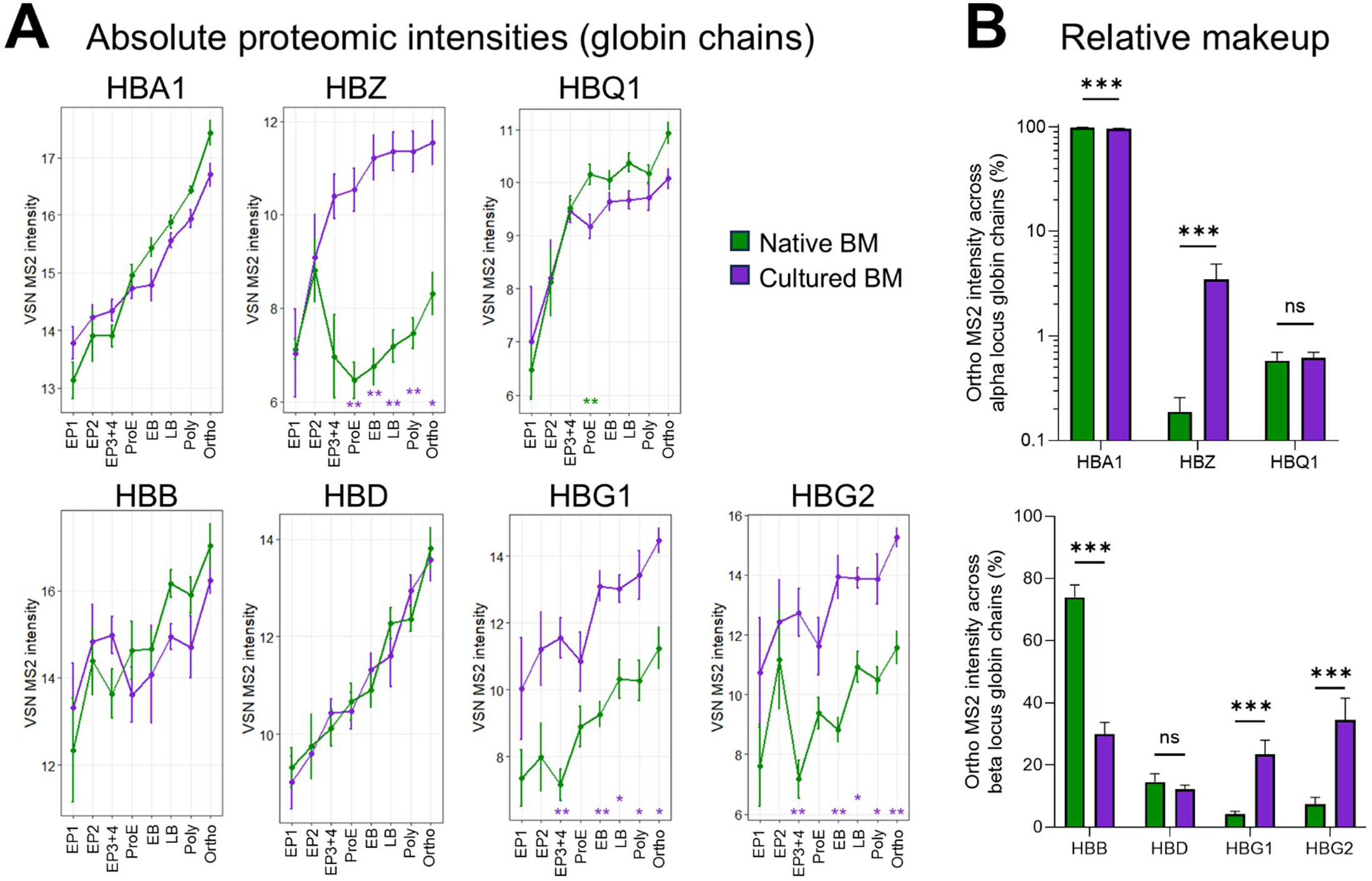
Absolute and relative quantification of globin chain proteins during erythropoiesis. **A)** VSN-MS2 intensities for detected globin chains across erythroid stages. Asterisks indicate statistical significance of differential protein expression. Green asterisks indicate significantly higher intensities in native bone marrow samples, purple asterisk indicate significantly higher intensities in culture-derived samples. ** = FDR < 0.01, * = FDR < 0.1. **B)** Frequency of globin chain MS2 intensities relative to total globin intensity. Asterisks indicate multiple comparison (Šidak)-corrected P-values. *** = P < 0.001, ns = not significant.

## Supplemental Methods

### Preparation of sorted cell samples for low-cell proteomics

The protocol utilized in this work was originally developed in Moura PL & Branca RM *et al*.^1^ and is further optimized in this study. Full details are provided below.

Evotips PURE (EvoSep) were prepared in advance to receive samples by i) washing with 20µl of solvent B (0.1% formic acid in acetonitrile), centrifuging (60s, 800 g); ii) soaking in isopropanol for 1min; and iii) equilibrating with 20µl of solvent A (0.1% formic acid in water), centrifuging (60s, 800 g) (2 times). The Evotips were then placed in an Evotip box containing 50mL of water to prevent drying.

Next, 5 µl of digestion buffer (0.2% DDM, 6%ACN, 100 mM TEAB pH8.5, 40 ng/µl trypsin (Rapizyme MS grade, Waters)) was added to each sample (in ∼4µl of FACS buffer) in the original tube used in FACS sorting (Protein Lo Bind, 1.5mL, Eppendorf) and mixed once by pipetting up and down. Samples were kept frozen on dry ice and only thawed immediately prior to addition of digestion buffer. The entire cell suspension mixed with digestion buffer (approx. 10 µl) was then transferred to a previously prepared Evotip. Evotips with samples were centrifuged for 3s and then placed Evotip box containing 50mL of water. The Evotip box was then incubated at 37°C for 4 hours with mild shaking. After incubation, 50µl of Solvent A was added to the Evotips to load the sample (centrifugation at 800 g, 60s). Followed by 20µl of Solvent A addition to the Evotips to rinse the sample (centrifugation at 800 g, 60s). Finally, 200µl of Solvent A was added onto Evotips to also prevent drying from the top, as recommended by the manufacturer’s instructions. Evotips were placed back in box with 50ml water.

Online LC-MS was performed using a Evosep One (Evosep, Odense, Denmark) coupled to a timsTOF SCP mass spectrometer (Bruker). The Evosep One LC system proceeded to inject sample onto the analytical column (Aurora Elite G3, C18, 15 cm long, 75 μm ID, 1.7 μm bead size, Ionoptics, Melbourne, Australia) constantly kept at 50°C by a heating oven (Column Toaster, Bruker, Bremen, Germany) using the Whisper 40 samples per day (40SPD) 31min gradient. The timsTOF SCP mass spectrometer operated with the CaptiveSpray source, capillary voltage 1500 V, dry gas flow of 3 L/min, dry gas temperature at 200°C. Collision energy was set as 20 eV for 1/k0 0.60 V.s/cm and 59 eV for 1/k0 1.60 V.s/cm2. Data was acquired using Timscontrol v 6.0.6 and Compass HyStar 6.3.1.8. The above conditions were used for data independent acquisition (DIA) diaPASEF (parallel accumulation-serial fragmentation). Precursor ions were selected in the range of 100-1700 m/z and mobility range 0.64-1.45 1/k0 IM. MS/MS was used with Bruker default dia-PASEF settings with 1 MS1 ramp, 8 MS/MS ramps, 24 MS/MS windows, mass range was 400–1000 m/z, and mobility range was 0.64–1.37 V.s/cm2. Parallel accumulation and serial fragmentation were performed with a cycle time of 0.95 s.

### Proteomics data analysis

Raw mass spectrometry data files were analyzed using Spectronaut vs. 19.4 (Biognosys) with the DirectDIA workflow for label-free quantification. Data search parameters included 1% FDR (false discovery rate) on both protein and peptide level, 2 as maximum missed cleavages, and Trypsin as enzyme type and cleavage rules. The search was performed against the Human Swissprot protein database (2024-01-10, 20413 canonical protein entries). The resulting MS2 protein intensity matrices from Spectronaut analysis were transformed using the variance-stabilizing normalization (VSN) function *vsn2* from the R package *vsn*,^2^ and batch-corrected per combined mass spectrometry and cell culture batch (*n* = 3 batches) using the *removeBatchEffect* function from *limma*.^3^ Proteins where insufficient detection was achieved were excluded from the dataset (detected in <50% of samples from either group under comparison). Following these initial steps, pairwise statistical comparisons of mass spectrometry data between culture and steady-state conditions were performed by pairwise comparison using Benjamini-Hochberg (BH) multiple comparison-corrected unpaired *t*-tests (if detection was sufficient in both groups) and Fisher’s exact tests (if detection was insufficient in one of the groups), after normalization of VSN intensity values against mean cell compartment VSN intensities. Time-course analysis of steady-state human erythroid differentiation was performed using the MaSigPro package^4^ and fitting erythroid transition between differentiation states to a time-course analysis (considering each differentiation state as a step in pseudotime). MaSigPro regression fitting was constrained to Q values < 0.10, calculated using the Benjamini-Hochberg method for false discovery rate-adjusted P-values. Following this, a backward stepwise regression fit was calculated at a significance level of alpha = 0.10 to identify the best regression model for each protein.

### Single-cell RNA sequencing data integration and analysis

Data matrices were obtained from Moura *et al*.^5^ (10X Chromium, RNA data from BM samples of 3 healthy individuals and 5 myelodysplastic syndromes [MDS] patients), Berastegui *et al*.^6^ (10X Chromium, RNA data from BM-derived control transduced CD34^+^ cells after 4 days of expansion and 14 days of erythroid culture conditions) and Triana *et al*.^7^ (BD Rhapsody, RNA and antibody-derived tag [ADT] data from BM samples of 6 healthy individuals).

Seurat v.5.4.0.^8^ was used to load, process and analyze the above data. Quality control filtering steps were applied to remove cells expressing less than 200 distinct genes, cells with a total percentage of mitochondrial reads above 15% and a total read count below 500 counts, as well as through removal of genes expressed in less than 3 cells. All samples were Log normalized and the 2000 most highly variable genes across all samples were identified and used as a basis for integration anchor identification and integration of both datasets. RNA data integration was anchored on a concurrently acquired batch of 2 healthy BM samples from Moura *et al*. After integration, the data were scaled and Principal Component Analysis (PCA) was performed. The 50 PCs with lowest P-values were selected for UMAP projection^9^ and clustering (through a shared nearest neighbor [SNN] modularity optimization-based clustering algorithm^10^ at a resolution parameter of 0.6). The dataset was filtered to include only early hematopoietic progenitors as well as terminally differentiating cells. After removal of undesired cell lineages, data were re-scaled and re-clustered at a resolution parameter of 8 (to improve specification ability within the relatively sparse early erythroid progenitors). ADT data from Triana *et al*. was overlaid on the dataset and, together with cluster expression data, utilized to manually annotate each cluster according to their proximal immunophenotypes and cluster-specific gene expression profiles, labelling in the erythropoietic continuum either to early progenitors HSCMPP, megakaryocyte-erythroid progenitors MEP, or the the immunophenotypes described within this study, namely EP1, EP2, EP3+4, ProE, EB, LB, Poly and Ortho. After labelling, further analyses were restricted to Chromium data from Moura *et al*. and Berastegui *et al*. Differential expression analyses were performed between sample groups through the Seurat-implemented non-parametric Wilcoxon rank sum test, with a log2 fold change threshold of 0.25 and a minimum group-wise percentage of DEG-expressing cells of 5%. The obtained P-values were adjusted with the Benjamini-Hochberg method and genes with an adjusted P-value below 0.001 in cluster analyses, or 10^-50^ in the total analysis of culture vs. BM, were reported as significantly differentially expressed.

## Supplemental tables

**Table S1:** Antibodies and fluorescent reagents.

| **Antibodies** |  |  |
| --- | --- | --- |
| CD3-PE-Cy5 | Biolegend | Cat no. 300410, RRID:AB_314064 |
| CD11b-PE-Cy5 | Biolegend | Cat no. 301308, RRID:AB_314160 |
| CD19-PE-Cy5 | Biolegend | Cat no. 302210, RRID:AB_314239 |
| CD56-PE-Cy5 | BD Biosciences | Cat no. 555517, RRID:AB_395907 |
| CD34-A700 | BD Biosciences | Cat no. 561440, RRID:AB_10715443 |
| CD105-Biotin | Biolegend | Cat no. 323214, RRID:AB_2293527 |
| Band 3-FITC | ARP American Research Products | Cat no. 08-0439-3, RRID:AB_3068600 |
| CD235a-PE | Biolegend | Cat no. 349106, RRID:AB_10641707 |
| CD71-APC-Cy7 | Biolegend | Cat no. 334109, RRID:AB_2563116 |
| CD49d-APC | Miltenyi Biotec | Cat no. 130-123-797, RRID:AB_2802060 |
| **Fluorescent reagents** |  |  |
| 7-AAD | eBiosciences | Cat no. 00-6993-50 |
| Hoechst 33342 | Sigma-Aldrich | Cat no. B2261 |
| DRAQ5 | Thermo Fisher | Cat no. 62251 |
| Streptavidin-BV650 | BD Biosciences | Cat no. 563855 |

